# A New Phylogenetic Framework for the Animal-adapted *Mycobacterium tuberculosis* Complex

**DOI:** 10.1101/384263

**Authors:** Daniela Brites, Chloe Loiseau, Fabrizio Menardo, Sonia Borrell, Maria Beatrice Boniotti, Robin Warren, Anzaan Dippenaar, Sven David Charles Parsons, Christian Beisel, Marcel A. Behr, Janet A Fyfe, Mireia Coscolla, Sebastien Gagneux

## Abstract

Tuberculosis (TB) affects humans and other animals and is caused by bacteria from the *Mycobacterium tuberculosis* complex (MTBC). Previous studies have shown that there are at least nine members of the MTBC infecting animals other than humans; these have also been referred to as ecotypes. However, the ecology and the evolution of these animal-adapted MTBC ecotypes are poorly understood. Here we screened 12,886 publicly available MTBC genomes and newly sequenced 17 animal-adapted MTBC strains, gathering a total of 529 genomes of animal-adapted MTBC strains. Phylogenomic and comparative analyses confirm that the animal-adapted MTBC members are paraphyletic with some members more closely related to the human-adapted *Mycobacterium africanum* Lineage 6 than to other animal-adapted strains. Furthermore, we identified four main animal-adapted MTBC clades that might correspond to four main host shifts; two of these clades are proposed to reflect independent cattle domestication events. Contrary to what would be expected from an obligate pathogen, MTBC nucleotide diversity was not positively correlated with host phylogenetic distances, suggesting that host tropism in the animal-adapted MTBC seems to be driven more by contact rates and demographic aspects of the host population rather than host relatedness. By combining phylogenomics with ecological data, we propose an evolutionary scenario in which the ancestor of Lineage 6 and all animal-adapted MTBC ecotypes was a generalist pathogen that subsequently adapted to different host species. This study provides a new phylogenetic framework to better understand the evolution of the different ecotypes of the MTBC and guide future work aimed at elucidating the molecular mechanisms underlying host specificity.

## Introduction

Tuberculosis (TB) remains a major concern both from a global health and economic point of view. With an estimated 10.4 million new human cases and 1.7 million fatalities every year, TB kills more people than any other infectious disease (WHO 2014). Moreover, bovine TB is responsible for an estimated US$3 billion annual economic loss in livestock production globally (Waters et al. 2012) and represents an ongoing threat for zoonotic TB in humans (Olea-Popelka et al. 2017). The causative agents of TB in humans and animals are a group of closely related acid-fast bacilli collectively known as the *Mycobacterium tuberculosis* complex (MTBC) (Brites et al. 2017, Malone et al. 2017). The human-adapted MTBC comprises five main phylogenetic lineages generally referred to as *Mycobacterium tuberculosis* sensu stricto (i.e. MTBC lineages 1-4 and lineage 7) and two lineages traditionally known as *Mycobacterium africanum* (i.e. MTBC lineages 5 and 6) (de Jong et al. 2010, Brites et al. 2017, Yeboah-Manu et al. 2017). Among the animal-adapted members of the MTBC, some primarily infect wild mammal species (Malone et al. 2017). These include *Mycobacterium microti* (a pathogen of voles) (Brodin et al. 2002), *Mycobacterium pinnipedii* (seals and sea lions) (Cousins et al. 2003), *Mycobacterium orygis* (antelopes) (van Ingen et al. 2012) and the “dassie bacillus” (rock hyrax) (Mostowy et al. 2004), which have been known for a long time, as well as the more recently discovered *Mycobacterium mungi* (mongooses) (Alexander et al. 2010), *Mycobacterium suricattae* (meerkats) (Parsons et al. 2013) and the “chimpanzee bacillus” (chimpanzees) (Coscolla et al. 2013). *Mycobacterium bovis* and *Mycobacterium caprae* on the other hand are mainly found in domesticated cattle and goats, but do also frequently spill over into many different wild animal species (Malone et al. 2017). *Mycobacterium canettii* is also considered part of the MTBC based on nucleotide identity; however *M. canettii* is likely an environmental microbe only occasionally causing opportunistic infections in humans (Koeck et al. 2010, Supply et al. 2013). We therefore use the term MTBC to refer to all the above mentioned members except *M. canettii*. Many of the names of the animal-adapted MTBC species were originally coined based on the animal they were first isolated from. For example, *M. orygis* was first identified in a captive oryx (van Soolingen et al. 1994) but has since then been isolated from many different host species including humans (van Ingen et al. 2012). Thus, the actual host range of *M. orygis* remains ill-defined (Malone et al. 2017). Similarly, for many of the animal-adapted members of the MTBC, only a few representatives have been isolated so far (e.g. only one in the case of the chimpanzee bacillus), limiting inferences with respect to the host range of these microbes. Moreover when studying host tropism, it is important to differentiate between maintenance hosts, in which the corresponding MTBC members traverse their full life cycle, including the transmission to secondary hosts, and spillover hosts, in which the infection leads to a dead end with no onward transmission (Malone et al. 2017). For example, *M. tuberculosis* sensu stricto is well adapted to transmit from human to human (Brites et al. 2015) and is occasionally isolated from cattle (Ameni et al. 2010). However *M. tuberculosis* sensu stricto is avirulent in cattle (Whelan et al. 2010, Villarreal-Ramos et al. 2018) and transmission from an animal back to humans is extremely rare (Murphree et al. 2011). Conversely, *M. bovis* is well adapted to transmit among cattle and does occasionally infect humans, mainly through the consumption of raw milk (Muller et al. 2013) or close contact with infected cattle, but transmission of *M. bovis* among immuno-competent humans is similarly uncommon (Blazquez et al. 1997).

The different members and phylogenetic lineages of the MTBC share a high nucleotide identity (>99.9%), and it has recently been suggested that they should be regarded as part of the same bacterial species (Riojas et al. 2018). The fact that these lineages also occupy different ecological niches, which is reflected in their host-specific tropism, supports a distinction into separate ecotypes (Smith et al. 2005). Yet, the host range of many of these animal-adapted MTBC members remain poorly defined, with respect to both maintenance and spillover hosts (Malone et al. 2017). In this study, we present and discuss a new phylogenetic framework based on whole genome sequences covering all known MTBC ecotypes. Based on this novel framework, we challenge previous assumptions regarding the evolutionary history of the MTBC as a whole, and point to new research directions for uncovering the molecular basis of host tropism in one of the most important bacterial pathogens.

## Methods

### MTBC genome dataset

We downloaded 12,886 genomes previously published and accessible from the sequence read archive (SRA) repository by December 2017 (Menardo et al. 2018). After mapping and calling of variants (see below), phylogenetic SNPs as in (Steiner et al. 2014) were used to classify genomes into human-adapted MTBC if they belonged to lineages 1 to 7 and if not, into non-human (hereafter referred to as “animal”) MTBC. All genomes determined as animal MTBC, as well as those classified as L5 or L6, were used for downstream analysis. We have furthermore newly sequenced four *M. orygis* genomes, two dassie bacillus genomes, eight *M. microti*, two *M. bovis* and one *M. caprae* (Supplementary Table 1). For downstream analysis, we selected the genomes published in (Comas et al. 2013) as representatives of other human MTBC, giving a total 851 genomes used in the downstream analysis (Supplementary Table 1).

### Bacterial culture, DNA extraction and whole-genome sequencing

The MTBC isolates were grown in 7H9-Tween 0.05% medium (BD) +/-40mM sodium pyruvate. We extracted genomic DNA after harvesting the bacterial cultures in the late exponential phase of growth using the CTAB method (Belisle et al. 1998). Sequencing libraries were prepared using NEXTERA XT DNA Preparation Kit (Illumina, San Diego, USA). Multiplexed libraries were paired-end sequenced on an Illumina HiSeq2500 instrument (Illumina, San Diego, USA) with 151 or 101 cycles at the Genomics Facility of the University of Basel. In the case of the *M. microti* isolates, DNA was obtained using the QIAamp DNA mini kit (Qiagen, Hilden, Germany) and libraries also prepared with the NEXTERA XT DNA Preparation Kit, were sequenced on an Illumina MiSeq using the Miseq Reagent Kit v2, 250-cycle paired-end run (Illumina, San Diego, USA).

### Bioinformatics analysis

#### Mapping and variant calling of Illumina reads

The obtained FASTQ files were processed with Trimmomatic v 0.33 (SLIDINGWINDOW: 5:20) (Bolger et al. 2014) to clip Illumina adaptors and trim low quality reads. Reads shorter than 20 bp were excluded from the downstream analysis. Overlapping paired-end reads were merged with SeqPrep v 1.2 (overlap size = 15) (https://github.com/jstjohn/SeqPrep") We used BWA v0.7.13 (mem algorithm) (Li et al. 2010) to align the reads to the reconstructed ancestral sequence of MTBC obtained as reported (Comas et al. 2010). Duplicated reads were marked by the Mark Duplicates module of Picard v 2.9.1 (https://github.com/broadinstitute/picard) and excluded. To avoid false positive calls, Pysam v 0.9.0 (https://github.com/pysam-developers/pysam) was used to exclude reads with alignment score lower than (0.93*read_length)-(read_length*4*0.07)), corresponding to more than 7 miss-matches per 100 bp. SNPs were called with Samtools v 1.2 mpileup (Li 2011) and VarScan v 2.4.1 (Koboldt et al. 2012) using the following thresholds: minimum mapping quality of 20, minimum base quality at a position of 20, minimum read depth at a position of 7x and without strand bias. Only SNPs considered to have reached fixation within an isolate were considered (at a within-host frequency of ≥90%). Conversely, when the SNP within-isolate frequency was ≤10% the ancestor state was called. Mixed infections or contaminations were discarded by excluding genomes with more than 1000 variable positions with within-host frequencies between 90% and 10% and genomes for which the number of within-host SNPs was higher than the number of fixed SNPs. Additionally, we excluded genomes with average coverage lower than 15x (after all the referred filtering steps). All SNPs were annotated using snpEff v4.11 (Cingolani et al. 2012), in accordance with the *M. tuberculosis* H37Rv reference annotation (NC_000962.3). SNPs falling in regions such as PPE and PE-PGRS, phages, insertion sequences and in regions with at least 50 bp identities to other regions in the genome were excluded from the analysis (Stucki et al. 2016). SNPs known to confer drug resistance as used in (Steiner et al. 2014) were also excluded from the analysis. Customized scripts were used to calculate mean coverageper gene corrected by the size of the gene. Gene deletions were determined as regions with no coverage to the reference genome. To identify deletions of regions and genes absent from the chromosome of H37Rv (e.g. RD900) unmapped reads resultant from the previous mapping procedure were mapped with reference to *M. canettii* (SRX002429) annotated using as reference NC_015848.1, following the same steps described above.

Phylogenetic reconstructionAll 851 selected genomes were used to produce an alignment containing only polymorphic sites. The alignment was used infer a Maximum likelihood phylogenetic tree using the MPI parallel version of RaxML (Stamatakis 2006). The model GTR implemented in RAxML was used, and 1,000 rapid bootstrap inferences followed by a thorough maximum-likelihood search (Stamatakis 2006) was performed in CIPRES (Miller et al. 2010). The best-scoring Maximum Likelihood topology is shown. The phylogeny was rooted using *M. canettii*. The topology was annotated using the package ggtree (Guangchuang et al. 2017) from R (Team 2018) and Adobe Illustrator CC. Taxa images were obtained from http://phylopic.org/. To remove redundancy and obtain a more even representation of the different MTBC groups for analysis of population structure and genetic diversity, we applied Treemer (Menardo et al. 2018) with the stop option *-RTL* 0.95, i.e. keeping 95% of the original tree length. The resulting reduced dataset was used for further analysis.

#### Population structure and genetic diversity

Population structure was evaluated using Principal Component Analysis (PCA) on SNP differences using the R package *adegent* (Jombart 2008). Genetic diversity was measured as raw pair-wise SNP differences for each MTBC lineage and ecotype if there were more than four genomes from a different geographic location, and as mean nucleotide diversity per site π using the R package *ape* (Paradis et al. 2004). π was calculated as the mean number of pair-wise mismatches among a set of sequences divided by the total length of queried genome base pairs which comprise the total length of the genome after excluding repetitive regions (see above) (Hartl et al. 2006). Confidence intervals for π were obtained by bootstrapping (1000 replicates) by re-sampling with replacement the nucleotide sites of the original alignments of polymorphic positions using the function *sample* in R (Team 2018). Lower and upper levels of confidence were obtained by calculating the 2.5th and the 97.5th quantiles of the π distribution obtained by bootstrapping.

## Results and Discussion

### Genome-based phylogeny reveals multiple animal-adapted clades

We assembled a total of 851 whole-genome sequences covering all known MTBC lineages and ecotypes. These included 834 genomes published previously, as well as four *M. orygis* genomes, two dassie bacillus genomes, eight *M. microti*, two *M. bovis* and one *M. caprae* newly sequenced here (Supplementary Table 1). We used a total of 56,195 variable single nucleotide positions extracted from these genome sequences to construct a phylogenetic tree rooted with *M. canettii*, the phylogenetically closest relative of the MTBC (Supply et al. 2013) (Figure 1). Our findings support the classification of the human-adapted MTBC into seven main phylogenetic lineages as previously reported (Gagneux et al. 2006, Gagneux et al. 2007, Firdessa et al. 2013). Classical genotyping studies and genomic deletion analyses indicated a single monophyletic clade for all the animal-adapted MTBC defined by clade-specific deletions in the Regions of Difference (RD) 7, 8, 9 and 10 (Brosch et al. 2002, Mostowy et al. 2002), and our new genome-based analysis confirms that all known animal-adapted members of the MTBC share a common ancestor at the branching point which is characterized by these deletions. Of note, the human-adapted MTBC Lineage 6 also shares this common ancestor, which has led to the hypothesis that Lineage 6 might have an unknown animal reservoir (Smith et al. 2006); however no such reservoir has yet been identified Yeboah-Manu et al. 2017) Due to the limitations of standard genotyping (Comas et al. 2009) and the limited phylogenetic resolution of RDs in the MTBC (Hershberg et al. 2008), previous classifications have considered all animal-adapted ecotypes as part of one phylogenetic clade, recently referred to as MTBC “Lineage 8” (Gonzalo-Asensio et al. 2014). However, our new genome-based data revealed that these animal-adapted ecotypes form separate animal-adapted clades, some of which are paraphyletic. For the purpose of this study, we discuss four of these animal-adapted clades which we named Clade A1 to A4.

**Figure 1.**
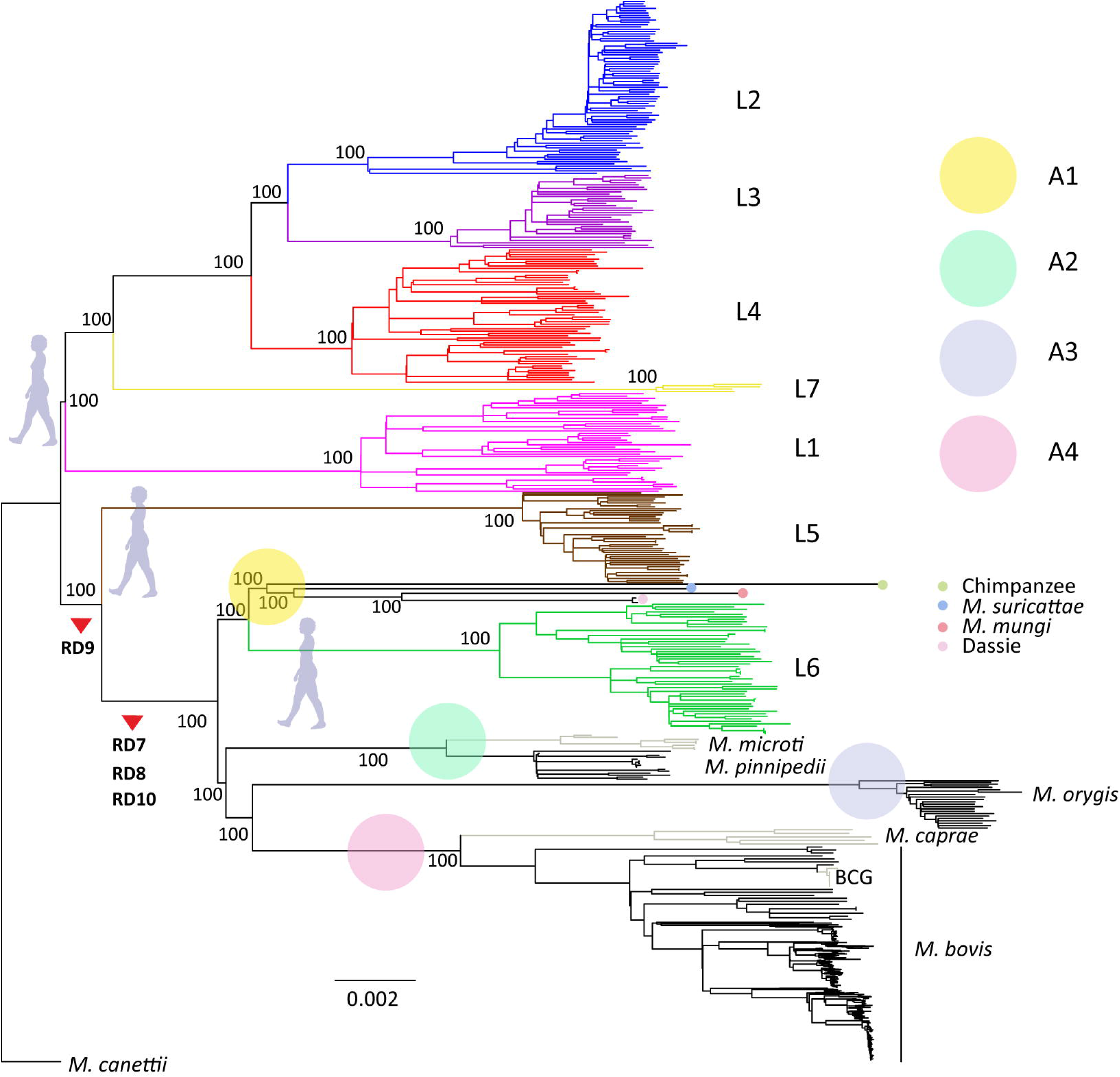
Maximum Likelihood topology of 322 human-adapted and 529 animal-adapted MTBC members. Branch lengths are proportional to nucleotide substitutions and the topology is rooted with *Mycobacterium canettii*. Support values correspond to bootstrap values. Main large deletions defining the animal-adapted MTBC are indicated by red arrows.

### The animal-adapted MTBC Clade A1

One important finding from our phylogenomic analysis was that *M. mungi, M. suricattae*, the dassie bacillus and the chimpanzee bacillus form a separate Clade A1, which clusters with the human-adapted MTBC Lineage 6 (Figure 2). Based on limited previous genotyping data (Huard et al. 2006), it was hypothesized that the dassie bacillus shared a common ancestor with *M. africanum* (i.e. MTBC Lineage 6) (Huard et al. 2006, Brites et al. 2015). Our new whole genome data now confirms this hypothesis, and at the same time, highlight the fact that Clade A1 is more closely related to the human-adapted Lineage 6 of the MTBC than to the other animal-adapted ecotypes. This observation has important implications for our understanding of the original emergence of the animal-adapted strains and the evolutionary history of the MTBC as a whole. Specifically, considering that Lineage 5 is human-adapted and basal to the RD7-10 defined lineages, according to the most parsimonious evolutionary scenario, the common ancestor defined by the deletions in RD7-10 was a human-adapted pathogen (Brosch et al. 2002, Mostowy et al. 2002), and given that MTBC Lineage 6 is human-adapted (de Jong et al. 2010, Yeboah-Manu et al. 2017), the jump into animal hosts had to occur at least twice. Alternatively, if this common ancestor was already animal-adapted, it had to jump back into humans during the emergence of Lineage 6. A slight modification of this latter scenario would see the RD7-10 common ancestor as a generalist capable of infecting and causing disease in multiple host species, which was followed by a host-specialization of the different ecotypes. The generalist notion could be further extended to the evolution of the MTBC as a whole. According to the most common view, the MTBC emerged as a human pathogen (Brosch et al. 2002, Mostowy et al. 2002, Smith et al. 2009). This notion is supported by the fact that except for the lineage defined by deletions in RD7-10, all other MTBC lineages are human-adapted (Brites et al. 2015). Moreover, all known *M. canettii* isolates have been obtained from human TB patients (Supply et al. 2013), suggesting that the common ancestor of all the MTBC was able to cause infections in humans. However according the latest available epidemiological data (Koeck et al. 2010), *M. canettii* and the other so-called smooth TB bacilli are most likely opportunistic pathogens with a reservoir in the environment (Supply et al. 2017). Hence, the ancestor of the MTBC could also have been a generalist initially, which then adapted to the various host species over time (Smith et al. 2009).

**Figure 2.**
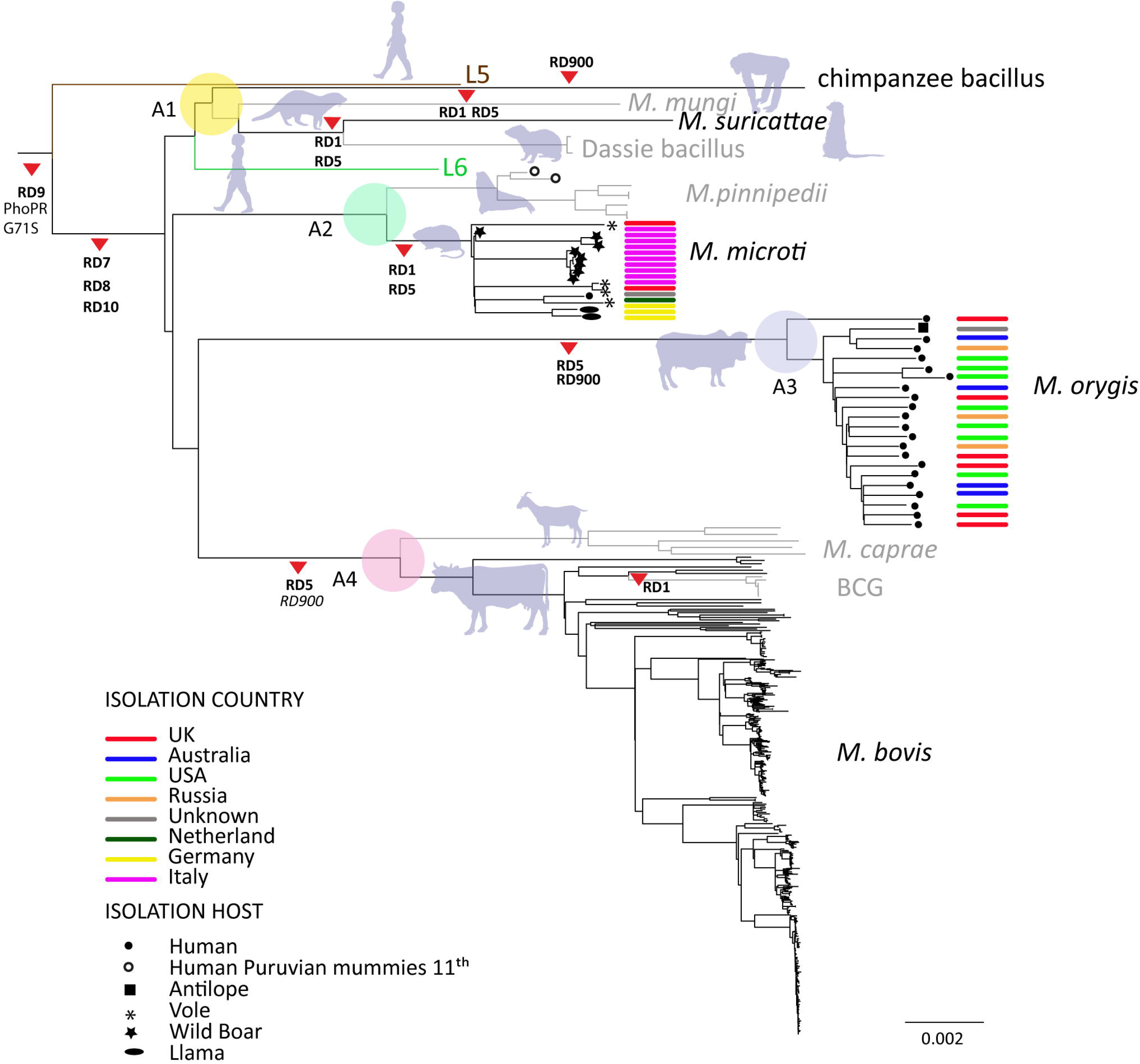
Topology showed in Figure 1 after collapsing all human-adapted branches. Branch lengths are proportional to nucleotide substitutions and the topology is rooted with *Mycobacterium canettii*. Main large deletions discussed in the text are indicated by red arrows. Those deletions which are polymorphic in terms of presence or absent within main clades are indicated in italics.

Another important characteristic of clade A1 is the absence of the region encoded by RD1 in *M. mungi, M. suricattae*, the dassie bacillus (Supplementary Table S2). RD1 encodes proteins that are essential virulence factors for MTBC in humans (further discussed below). Our data confirm that *M. mungi, M. suricattae*, the dassie bacillus all have deleted the region corresponding to RD1. This deletion is not present in the chimp bacillus suggesting that RD1 might be essential for virulence in primates as proposed previously (Dippenaar et al. 2015).

### The animal-adapted MTBC Clade A2

Similar to Clade A1 that comprises pathogens adapted to wild animals, Clade A2 consists of two ecotypes mainly affecting wild animals, namely *M. microti* and *M. pinnipedii*. In addition, Clade A2 also includes MTBC genomes isolated from pre-Columbian human remains published previously (Bos et al. 2014). These ancient genomes are most closely related to *M. pinnipedii*, suggesting possible cases of zoonotic TB transmission resulting from the handling and consumption of seal or sea lion meat at the time. Contemporary *M. pinnipedii* is known to infect humans occasionally (e.g. zoo keepers or seal trainers), but no human-to-human transmission has been documented to date. *M. microti* was originally isolated from voles in the 1930s (Wells 1937), but has since then been found in cats, pigs, llamas and immune-compromised humans (Brodin et al. 2002, Frota et al. 2004, Smith et al. 2009). Here we report 8 new *M. microti* genomes isolated from wild boar. Based on the 15 *M. microti* genomes included in this analysis, some host-specificity of particular sub-groups with this ecotype might be suggested, but analysis of a larger sample is needed to explore this possibility further. To our knowledge, *M. microti* has not been reported outside Europe, as infections in llamas pertain to captive animals in Europe (Oevermann et al. 2004) and represent probable spillovers from other hosts. Furthermore, the *M. microti–like* strain isolated from a rock hyrax has been likely misclassified (Clarke et al. 2016). Many of the animals species infected by *M. microti* occur across Eurasia which might therefore also correspond to the geographic range of *M. microti*. One of the important characteristics of all *M. microti* strains is the deletion of RD1 (Brodin et al. 2002), which is independent of the one described for the some of the members of Clade A1, and which is the most important virulence attenuating mutation in the *M. bovis* BCG vaccine (Pym et al. 2002). In support of the low virulence of *M. microti* in humans, and in contrast to *M. bovis* and *M. orygis* (see below), we detected only one infection with *M. microti* (from an immune-compromised patient (van Soolingen et al. 1998) among all the human isolates queried in the public domain (see methods).

### The animal-adapted MTBC Clade A3

In contrast to the animal Clades A1 and A2 that include multiple MTBC ecotypes infecting various wild animal host species, A3 comprises only genomes belonging to *M. orygis*. Even though *M. orygis* has been isolated from many different wild and domestic animals (Dawson et al. 2012, Gey van Pittius et al. 2012, van Ingen et al. 2012, Thapa et al. 2015, Thapa et al. 2016, Rahim et al. 2017), a large proportion of isolates reported to date are actually from human TB patients. One of the first detailed studies reporting on the genotypic properties of *M. orygis* strains included a total of 22 isolates, 11 of which originated from humans (van Ingen et al. 2012). The majority of the remaining isolates came from various zoo animals from the Netherlands and South Africa, which included three waterbucks, two antelopes, one deer and one oryx. A recent study from New York reported whole genome data from eight *M. orygis* isolates from human patients (Marcos et al. 2017). Another recent report from Birmingham, UK identified 24 *M. orygis* among 3,128 routinely collected human MTBC isolates (Lipworth et al. 2017). Similarly, eight *M. orygis* isolates were reported among 1,763 human TB cases from Victoria, Australia (Lavender et al. 2013), the genomes of four of which are newly reported here (Figure 1 and Figure 2). Importantly, all human *M. orygis* isolates, for which the relevant information was reported, were from patients born in India, Pakistan, Nepal or “South Asia”, except for one with a reported origin in “South East Asia” (Dawson et al. 2012, van Ingen et al. 2012, Lavender et al. 2013, Marcos et al. 2017). This also includes one patient who immigrated from India to New Zealand and infected a dairy cow there (Dawson et al. 2012). One recent study reported 18 *M. orygis* isolates from dairy cattle in Bangladesh (Rahim et al. 2017). These isolates grouped into three distinct MIRU-VNTR clusters, with the largest cluster including two additional *M. orygis* isolates from captive monkeys. The authors propose that *M. orygis* is endemic among wild and domestic animals across South Asia and thus of relevant One Health significance. Based on the available evidence summarized above, and given that *M. orygis* shares a common ancestor with Clade 4 (Figure 1), which comprises *M. bovis* and *M. caprae* primarily adapted to domestic animals (further discussed below), we extend this notion and hypothesize that *M. orygis* is primarily a pathogen of cattle in South Asia, leading to zoonotic TB in humans through e.g. the consumption of raw milk. This scenario is the most parsimonious explanation for why *M. orygis* has repeatedly been isolated from South Asian migrants living in low TB-endemic countries in Europe, USA and Australia (Dawson et al. 2012, van Ingen et al. 2012, Lavender et al. 2013, Marcos et al. 2017). The genetic distance among the *M. orygis* identified in this study also supports this scenario, as the genomes of these isolates differ on average by 231 SNPs, suggesting independent infections in their countries of origin (Figure 3). Broader in-depth molecular analyses of cattle TB in South Asia, for which little data currently exist despite it representing a major public health threat (Rahim et al. 2017, Srinivasan et al. 2018) are needed to verify our hypothesis. Regarding *M. orygis* reported in animals other than cattle, our hypothesis would suggest that these likely represent spillovers from infected cattle, similar to the situation seen in *M. bovis* (Malone et al. 2017). In support of this view, except for one case isolated from a free-ranging rhinoceros in Nepal (Thapa et al. 2016), all *M. orygis* reported in un-domesticated animals were associated with zoos, farms or other forms of captivity where these wild animals might have come into contact with *M. orygis* infected cattle or humans (Gey van Pittius et al. 2012, van Ingen et al. 2012, Thapa et al. 2015, Rahim et al. 2017).

**Figure 3.**
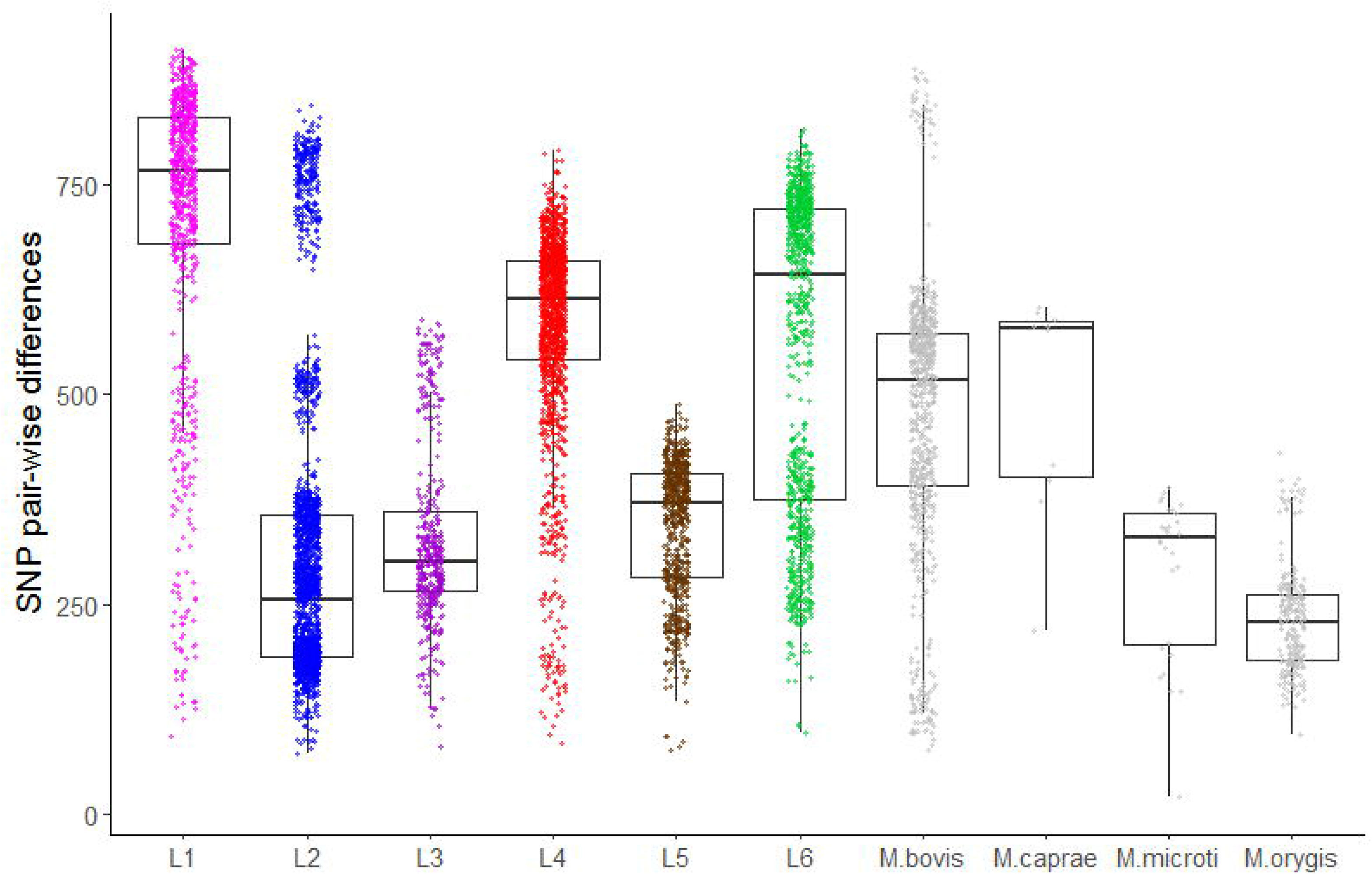
Pair-wise SNP distances within lineage and ecotype from human and animal-adapted MTBC, respectively. Each box corresponds to the 25% and 75% quantiles, the black line represents the median and the whiskers extend to 1.5 times the interquartile range.

### The animal-adapted MTBC Clade A4

Clade A4 includes the classical members of the animal-adapted MTBC, i.e. *M. bovis, M. caprae* and all the *M. bovis* BCG vaccine strains (Figure 2). Much work has been done on the genetic characterization of these MTBC members (Mostowy et al. 2005, Huard et al. 2006, Smith et al. 2006, Muller et al. 2009, Copin et al. 2014, Malone et al. 2017), and thus we will not discuss these in any further details here. One exception is the deletion RD900, which has been described as a region specific to L6 and for which, presence and absence in *M. bovis* has been disputed (Bentley et al. 2012, Malone et al. 2017). The results of mapping with respect to *M. canettii* reads which remained unmapped to the chromosome of H37Rv, revealed that RD900 is polymorphic within *M. bovis*, within BCG strains and within *M. caprae*. In contrast, the region encoded by RD900 is deleted in all *M. orygis* genomes analyzed (Figure 2).

We end this section by speculating that if our hypothesis regarding the host range of *M. orygis* is true, Clade A3 and Clade A4 might reflect the two independent cattle domestication events known to have occurred in the Fertile Crescent and Indus Valley, respectively (Loftus et al. 1994). The corresponding domesticated forms emerging form the ancestral aurochs *(Bos primigenious)* are the sub-species *Bos taurus* and *Bos indicus*. Hence, *M. bovis* might have adapted to *B. taurus* whereas *M. orygis* might be better adapted to *B. indicus*. While highly speculative at this stage, this hypothesis could be tested experimentally (Villarreal-Ramos et al. 2018).

### MTBC genetic diversity and host specificity

From an ecological perspective, pathogen diversity is generally positively correlated with host diversity especially in the case of obligate pathogens (Kamiya et al. 2014). Given the broad MTBC range of hosts, we explored how the genetic diversity is [partioned](http://www.thesaurus.com/browse/partitioned) within the MTBC and if the genetic diversity of the animal-adapted MTBC was higher than that of the human-adapted MTBC. To obtain a more balanced representation of the different MTBC groups and remove redundancy caused by an over-representation of very closely related isolates which tell us little about macro-evolutionary processes, we used Treemer (Menardo et al. 2018) and reduced our dataset from 851 to 367 genomes while keeping 95% of the original total tree length. We performed principal component analysis (PCA) on the matrix of SNP distances correspondent to the non-redundant data set (n=367) (Figure 4). The resultant groups correspond largely to the results obtained with the phylogenenetic approach. The first principal component (PC1) explains 20.5% of the variation in genetic differences and highlights the contrast between “modern” human MTBC lineages (Lineages 2, 3 and 4) and Lineages 1, 5 and 7, which on their own formed very distinct groups. Lineage 6 appears closer to the animal MTBC but separated from clade A1. Interestingly, despite a clear separation between the human-adapted and animal-adapted MTBC (except for Lineage 6), PC1 contrasts more prominently the different human-adapted lineages than the different animal-adapted ecotypes (Figure 4).

**Figure 4.**
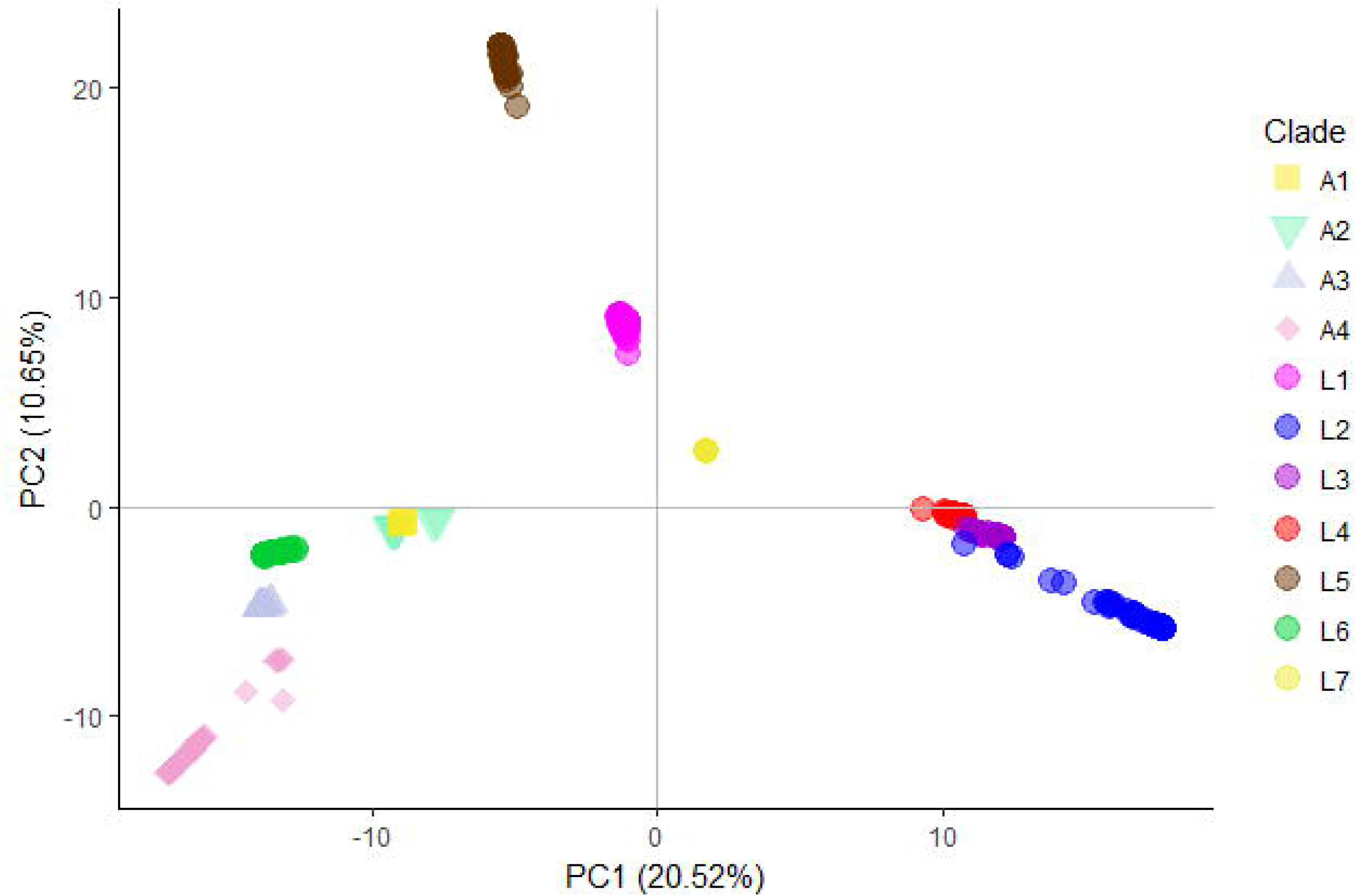
Principal Component Analysis (PCA) derived from whole-genome SNPs. The two first principal components are shown.

As a measure of genetic diversity, we estimated the mean nucleotide diversity per site (π) of human versus animals isolates. The estimates indicate that two randomly picked human isolates differ on average by 0.0345% nucleotide differences (95% CI: 0.0337%-0.0352%) whereas animal isolates differ on average by 0.0313% (95% CI: 0.0305%-0.0321%). Despite non-overlapping confidence intervals, the difference between both π estimates is very small (0.003%) indicating that the genetic diversity which has emerged within animal and human MTBC is of similar magnitude. The estimates of π reflect both the diversity within each lineage/ecotype, as well as the diversity between lineages/ecotypes, resulting from older evolutionary events leading to the emergence of the latter. Whereas our sampling of the human MTBC reflects both pre- and post-lineage diversification reasonably well, the animal MTBC samples are most likely a poor representation of the genetic diversity resulting from diversification processes within each ecotype, with the possible exception of *M. bovis* (Figure 3) We thus compared the raw SNP differences among one random representative of each human and animal-adapted MTBC lineage and ecotype (Figure 5). The SNP differences accumulated in the different human-adapted lineages can be as high, or even higher than the genetic differences that separate MTBC strains infecting a broad taxonomic range of mammal species other than humans. Thus host-specificity in the MTBC cannot be easily explained only by quantitative genetic differences among the different animal-adapted MTBC ecotypes. In the light of the fact that in the MTBC, as in other bacteria, genomic variants caused by large deletions are pervasive (Bolotin et al. 2015) and genomes evolve towards a reduction of gene content as no horizontal gene transfer has been found in extant populations of the MTBC, it is also unlikely that the acquisition of new genes underlies host specificity. In support of this, after mapping reads using *M. canettii* as a reference, we found no regions that would be present in all representatives of each the different animal ecotype genomes and absent from human-adapted MTBC genomes. Several genomic deletions have been described in the genomes of animal-adapted MTBC members which we could also confirm here (Supplementary Table 2). Some of those deletions, e.g. RD1 and RD5, have been shown to impact virulence in different ways (Lewis et al. 2003, Dippenaar et al. 2015, Ates et al.(2018). In the case of RD1 and RD5, the deletion events seem to have occurred independently in different animal MTBC ecotypes (Figure 2) suggesting that the former have provided a fitness gain and were involved in the adaptation to new hosts (Brodin et al. 2002, Dippenaar et al. 2015, Ates et al. 2018). However, RD5 has also independently evolved and shown to impact virulence in the human adapted L2 Beijing sub-lineage (Ates et al. 2018). Taken together, this suggests that MTBC genomes are extremely robust in terms of host adaptation, and that interactions between different genes in the different ecotypes will be key determinants of host specificity in the MTBC.

**Figure 5.**
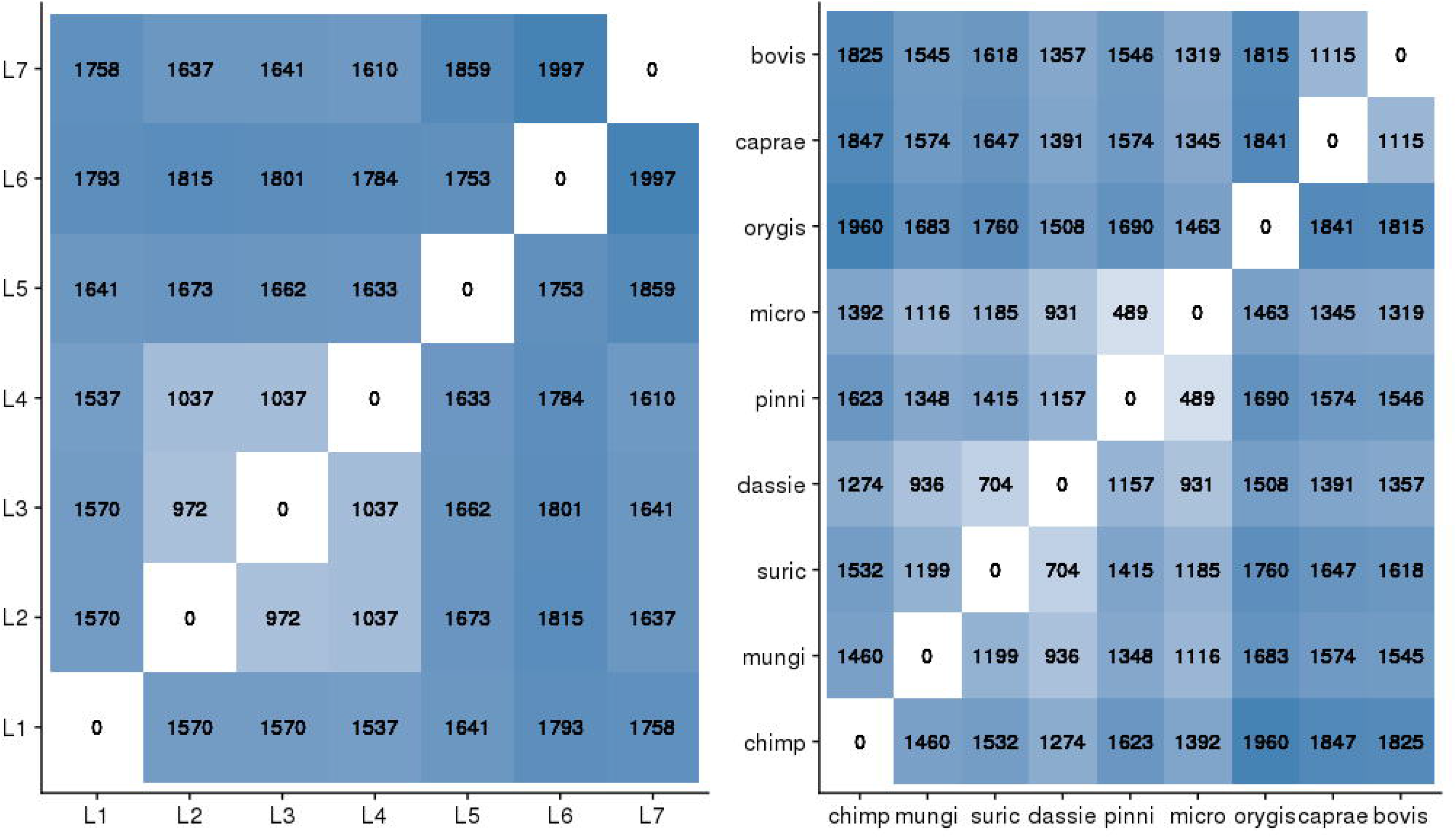
Pair-wise SNP distances between one randomly chosen representative of each human adapted MTBC lineage A) and animal-adapted MTBC B).

### Evolutionary scenarios for the evolution of the animal-adapted MTBC

The different MTBC members have adapted to infect a broad range of mammalian species, ranging from micro-mammals with short life-spans to humans, indicating that host shifts to distantly related hosts have occurred throughout the evolution of the MTBC. However, these host shifts have not emerged from any random phylogenetic branch of the MTBC as most of the human-adapted MTBC lineages are monophyletic and possibly locally adapted to different human populations (Fenner et al. 2013, Gagneux 2018). Host range expansion seems to have occurred after the split between Lineage 5 and the ancestor of Lineage 6 and all the animal ecotypes. The most parsimonious explanation is thus that the ancestor pathogen of the extant animal-adapted MTBC and Lineage 6 was a generalist with the ability to cause infections in many different kinds of hosts. A series of genetic events have been put forward by (Gonzalo-Asensio et al. 2014) to explain the decreased virulence of *M. africanum* L5 and L6 and the animal MTBC members compared to *M. tuberculosis* sensu stricto. A nonsynonymous mutation on the codon 71 in the *phoR* gene (Figure 2) which has emerged in the common ancestor of *M. africanum* L5 and L6 and of the animal-adapted strains, if transferred to a *M. tuberculosis* sensu stricto background leads to decreased virulence in mice and primary macrophages (Gonzalo-Asensio et al. 2014). This decrease in virulence is mediated by a decrease in the secretion of ESAT-6 which among other virulence factors is regulated by*phoPR* genes. The work of (Gonzalo-Asensio et al. 2014) shows that in L6, the loss of virulence was compensated by the RD8 deletion which restored the secretion of ESAT-6 independently of *PhoPR*. RD8 is common to L6 and all the animal ecotypes (but not L5, Figure 2), thus how the effects of *PhoPR* are restored in L5 remains unknown. This and related events could be at the origin of a putative generalist pathogen with compromised virulence in its original human host, and for which infecting other hosts represented fitness gains leading to the host range expansion we see today.

Based on the known geographic ranges of the animal-adapted MTBC ecotypes, we suggest two main divisions after the emergence of the ancestor of L6 and the animal ecotypes (AncL6-_A_, Figure 6); A series of specialization events which have occurred within Africa leading to the emergence of L6 in humans and clade A1 in several wild mammal species. With the exception of the chimp bacillus, these ecotypes have all been sampled in Southern Africa (Clarke et al. 2016). However, the extant geographic distributions of the hosts are not restricted to Southern Africa (except for Meerkats), additionally they have several overlapping areas and as a whole, form a continuum ranging from West-Africa to Southern-Africa (see http://www.iucnredlist.org/). Another series of specialization events might have happened outside Africa as suggested by the extant distribution of *M. orygis* and *M. microti* (Figure 6). Given that the maintenance hosts of strains that comprise A3 and A4 are domesticated species, one possible scenario is that the ancestor of Anc_A2-A3-A4_ was carried by human populations as they migrated from Africa to the rest of the world (Figure 6). This ancestor could have been transferred posteriorly to different cattle and other livestock species which were domesticated outside Africa and independently in different parts of world as suggested in the discussion of clade A3 above, and become extinct in human populations. The example of the three human Peruvian mummies circa 1000 years old, which were infected with what is known today as *M. pinnipedii* (Bos et al. 2014), despite the difficulties in defining if humans were maintenance or spill-over hosts, illustrate the plausibility of such a scenario. Alternatively, Anc_A2-A3-A4_ might have been brought outside Africa by another migratory species with close contact to livestock. The jump from the ancestor Anc_A2-A3-A4_ to clade A2, which comprises such different host species, is not easily explained without invoking an environmental reservoir. This cannot be excluded as *M. bovis* and *M. microti* can possibly survive in the environment (Courtenay et al. 2006, Kipar et al. 2014).

**Figure 6.**
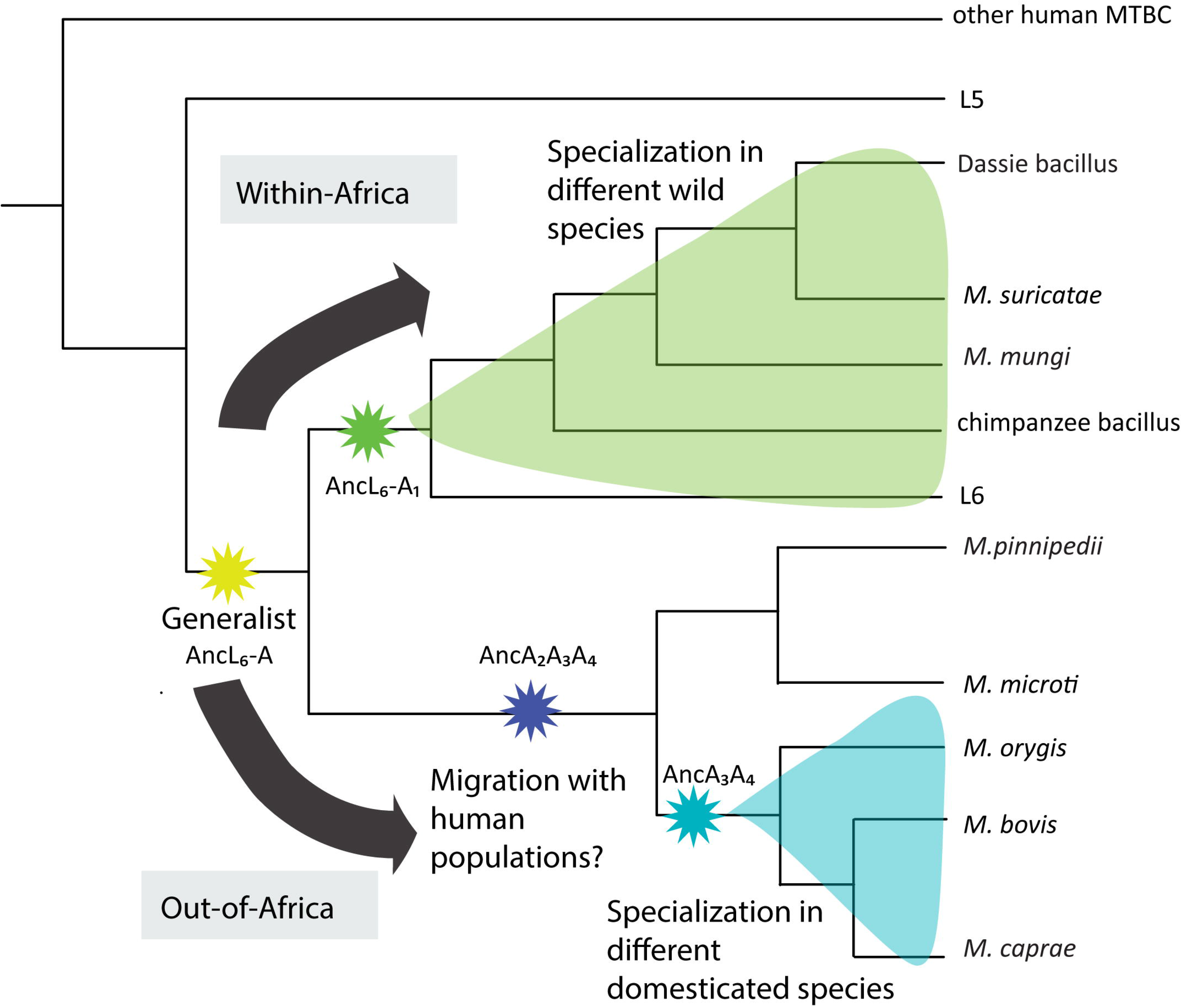
Schematic illustration of the putative evolutionary history of the animal-adapted MTBC. The length of the branch is not proportional to genetic distances.

The biology of pathogen jumps into new hosts involves three main steps (Woolhouse et al. 2005); i) exposure of the pathogen to a new environment, i.e. contact rates between hosts or between hosts and an environmental reservoir, ii) the ability to infect the new host, which most commonly decreases with the genetic distance from the ancestral host, and iii) transmissibility within the new host population. Generally, when the complete host ranges and the known geographic distributions are taken into account in the animal-adapted MTBC ecotypes, geographic proximity between hosts and therefore contact rates seemed to have played a more important role in determining host range and specialization than the phylogenetic distance among hosts. A corollary of these considerations and given that one important contributor to ii) is the ability to avoid or supress the host immune system, is that the immune repertoire of the host may have played a less important role in determining the host range of the different animal MTBC ecotypes compared to i) or iii) as long as the hosts were mammalian species. There are exceptions to this, e.g within Clade A1 moongoses and meerkats belong to the same taxonomic family (Clarke et al. 2016). However, in this case, host geographic range, ecology and phylogenetic distances are not independent, blurring conclusions. One important characteristic common to all host species in which the different MTBC members cause sustainable infections is that they attain high population densities, even if predominantly seasonally as in the case of pinnipeds (Cassini 1999). This characteristic might have been one of the most important determinants in the evolution of the different MTBC ecotypes and in particular, of their mode of transmission. Whereas the ability to cause pulmonary infections is essential for transmission among humans, in other animals, routes of infection other than aerosol transmission seem to play an important role, e.g. grazing contaminated pasture leads probably to a significant proportion of infections by *M. bovis* in cattle (Phillips et al. 2003), *M. mungi* can transmit directly through abrasions resultant from foraging activity of banded mongoose (Alexander et al. 2010, Malone et al. 2017), and transmission through skin lesions in *M. microti* has also been suggested (Kipar et al. 2014).

### Concluding remarks

There are several reports about animal-adapted members of the MTBC infecting humans, wild and domestic animals, but an overarching analysis of all information available is required. In this study, we have combined all available information about animal-adapted MTBC strains and expanded it by sequencing more animal-adapted MTBC strains gathering the most comprehensive whole genome dataset of animal-adapted MTBC to date. We have used genomic analysis to elucidate the evolutionary history of the animal-adapted MTBC and have confirmed that the former are paraphyletic and that at least four different main clades can be defined. The phylogeny presented together with the known host range would be compatible with two realistic scenarios during the evolutionary history of the non-human MTBC, both involving more than one host jump. One scenario would present the ancestor of the group including L6 and all animal-adapted clades as a generalist capable of infecting a wide group of mammals, and different host adaptations would have occurred thereafter. An alternative scenario proposes that the ancestor of L6 and animal-adapted MTBC was adapted to humans, and subsequent host jumps lead to the host specificity of the four clades.

We found no correlation between genetic diversity of the pathogen and the phylogenetic distance of the host, as animal-adapted MTBC strains are not more diverse in average than human-adapted strains. Based on the current known host-ranges and geography of the animal-adapted MTBC, we propose that host expansion has been driven to a great extent by host geographical proximity, i.e. by contact rates among different species of mammals, and by high host population densities rather than by host genetic relatedness.

## Author contributions

DB, CL, MC and FM have analysed the data. DB, SG and MC wrote the manuscript. BB, RW, AD, SP, MB, CB, SB, and JF contributed reagents and performed the experiments.

## Acknowledgements

Calculations were performed at sciCORE (http://scicore.unibas.ch/) scientific computing core facility at University of Basel. Library preparation and sequencing was carried out in the Genomics Facility Basel.

## Supplementary Material

SupplementaryTable1.xls

SupplementaryTable2.xls

